# Decoding the role of DNA sequence on protein-DNA co-condensation

**DOI:** 10.1101/2024.02.24.581870

**Authors:** Rohit Kumar Singh, Pinaki Swain, Mahipal Ganji, Sandeep Choubey

**Affiliations:** The Institute of Mathematical Sciences, CIT Campus, Tharamani, Chennai 600113, India; Homi Bhabha National Institute, Training School Complex, Anushaktinagar, Mumbai 400094, India; Department of Biochemistry, Indian Institute of Science, Bangalore, India

## Abstract

The eukaryotic genome is organized within the cell nucleus through three-dimensional compaction. The physical principles that govern genome organization *in vivo* remain less understood. Phase separation of protein and DNA has emerged as an attractive mechanism for reshaping chromatin and compacting the genome. *In vitro* studies have shed light on the biophysical principles of protein-DNA condensates driven by protein-protein and protein-DNA interactions. However, the role of DNA sequence and its impact on protein-DNA condensation remains elusive. Guided by experiments, this paper presents a simple polymer-based model of protein-mediated DNA condensation that explicitly incorporates the influence of DNA sequence on protein binding. Using coarse-grained Brownian dynamics simulations, we demonstrate that, in the case of a homogeneous DNA, only one condensate forms in equilibrium. In sharp contrast, DNA sequence heterogeneity can result in the coexistence of multiple condensates, giving rise to the formation of structures resembling pearl-necklaces. Interestingly, we observe that protein binding affinity of interfacial DNA governs the capillary forces arising from the protein-DNA condensates. To demonstrate the usefulness of our modeling framework, we compare the simulation results against published data for co-condensation of Dps, Sox2, and HP1. We find that while Dps exhibits sequence-independent binding, DNA sequence heterogeneity dictates the co-condensation of Sox2 and HP1 with DNA. Overall, the framework developed here can be harnessed to gain mechanistic insights into the role of DNA sequence on protein-DNA co-condensation and pave the way for developing a deeper understanding of genome organisation.

## 1 Introduction

The eukaryotic genome is spatially organized within the cell nucleus through three-dimensional (3D) compaction. Such spatial organizational architecture of the genome encodes for the spatio-temporal regulation of gene expression, DNA replication etc [1–4]. The genome is organized hierarchically, exhibiting distinct yet ubiquitous architectural features at different length scales such as the loops, domains, and compartments that the chromatin forms[4,5]. Unfurling the physical principles that regulate and remodel chromatin *in vivo* remains a central objective of regulatory biology.

Recent *in vitro* experiments suggest that transcription factors and DNA can undergo co-condensation through the collective behavior of thousands of proteins and DNA, driven by protein-protein and protein-DNA interactions [6–9]. The resulting protein-DNA co-condensate ensnares a specific amount of DNA, exerting force on the surrounding free DNA, known as capillary force[9–12]. Capillary forces, arising from the interfacial physics of protein-DNA condensates, have the potential to serve as a mechanism for regulating and reshaping chromatin. This capability holds promise in influencing the 3D architecture of the genome.

Experimental studies have investigated the interfacial physics of protein-DNA co-condensates and their impact on capillary forces by employing optical tweezers and coverslip-based assays, as shown in Figure 1 A, B. For instance, Quail et al. utilized total internal reflection fluorescence (TIRF) microscopy to explore the co-condensation of *λ*-phage DNA and the pioneer transcription factor protein FoxA1 [10]. Their findings revealed that the FoxA1-DNA co-condensate could generate forces on the order of 0.2 piconewton (pN). Another study highlighted that PARP1-DNA co-condensates generate capillary forces in the sub-piconewton range [11]. Intriguingly, a more recent investigation demonstrated that the pluripotent factor Sox2 has the capability to generate up to 7 pN of capillary forces [9]. While a combination of theory and experiments have provided valuable physical insights into the formation of protein-DNA co-condensates, a comprehensive understanding of the interfacial physics of co-condensates and their impact on the resultant capillary forces remains in its infancy. Specifically, the influence of DNA sequence heterogeneity on protein-DNA co-condensation and capillary forces remains less explored. The objective of this manuscript is to establish a modeling framework that would enable the study of the physics of protein-DNA co-condensates in the presence of DNA sequence heterogeneity.

For this purpose, we have employed a simple model of protein-DNA co-condensation that explicitly incorporates DNA sequence heterogeneity for protein binding. Moreover, our model is specifically designed to capture two types of *in vitro* systems used in the study of protein-DNA co-condensation: tweezer and coverslip-based assays. Using coarse-grained Brownian dynamics simulations, we demonstrate that, in the case of a homogeneous DNA polymer, only one condensate forms in equilibrium. In contrast, contingent upon DNA sequence heterogeneity, multiple condensates of varying sizes can coexist on the same DNA at equilibrium. Our study uncovers that the protein binding affinity of interfacial DNA dictates the magnitude of capillary forces. This observation offers a mechanism for regulating global forces on the genome through alterations at the DNA base pair level. In summary, our modeling framework offers a valuable tool for leveraging existing *in vitro* systems to gain mechanistic insights into the biophysics of protein-DNA co-condensation.

**Figure 1:**
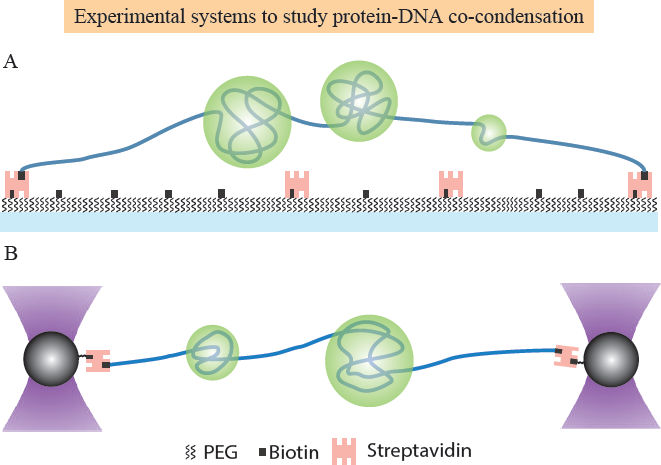
Experimental systems to study protein-DNA co-condensation. **A** Schematic of experimental setup wherein the ends of a single DNA molecule (blue) are tethered to the glass slide using biotin-streptavidin interactions. Proteins (green) phase-separate with DNA to form protein-DNA co-condensates. **B** Schematic of the optical tweezers assay: a DNA molecule (blue) is held between two optically trapped beads (black) via biotin–streptavidin interactions.

## 2 Model

To explore the impact of DNA sequence on the co-condensation of protein-DNA complexes and the subsequent generation of capillary forces, we propose a simple polymer-based model for protein-DNA binding. This model explicitly considers both protein-protein interactions and sequence-dependent protein-DNA interactions.

Informed by *in vitro* studies, as shown in Figure 1, we consider a 5 kb-long DNA as a semiflexible polymer consisting of 500 monomers, tethered at both ends within a cuboidal box. In accordance with previous studies [13], the persistence length of DNA is considered to be 150 bps, which translates to 15 monomers in our model. The DNA is in contact with proteins. Without any loss of generality, we assume that each monomer corresponds to a stretch of 10 bases that are occupied by a bound protein molecule. Indeed various prokaryotic and eukaryotic proteins occupy 10 bases on the DNA [14,15]. We assume that the proteins are spherical beads with dimensions identical to the monomers. The sequence heterogeneity of DNA is represented by distinct arrangements of monomers along the DNA contour, where each monomer exhibits varying binding affinity for the proteins. To dissect the impact of DNA sequence on protein-DNA co-condensation, first, we introduce a null model of homogeneous DNA. In this model, each monomer has an identical binding affinity of 2 *k_B_T* for proteins (see Figure. 2A)[7]. In contrast, we examine three distinct heterogeneous models for the DNA. 1) Heterogeneous DNA I, in this model, the DNA is represented as an ABA block copolymer based on differential attractive interaction with proteins. DNA consists of a high-affinity region spanning 250 monomers at the center, with a protein-DNA binding affinity of 2.25 *k_B_T*. This high-affinity region is surrounded by low-affinity regions on either side, each containing 125 monomers, with a protein-DNA binding affinity of 1.75 *k_B_T*. This ensures that the average protein-DNA binding affinity remains identical to the homogeneous case (see Figure. 2B). 2) Heterogeneous DNA II, in this model, five blocks of high-affinity regions of 2.25 *k_B_T*, each spanning 50 monomers along DNA contour, are interspersed with blocks of low-affinity regions with 1.75 *k_B_T* and span 50 monomers. The average protein-DNA binding affinity for this model is identical to the homogeneous case (see Figure. 4A). 3) Partial *λ*-DNA, to model a biologically realistic DNA, we consider the last 5 kb of the lambda DNA (see Figure. 5A and Methods section). For simplicity, we assume that each protein has a preference for binding to AT-rich regions. Indeed, there exist numerous proteins, in bacteria and higher organisms, that broadly bind AT or GC-rich sequences [16–18]. We assign a corresponding interaction energy to each monomer based on the AT content of each 10-base sequence (see the Methods and SI for details).

Following the experiments, we tune the normalized end-to-end distance of DNA and the bulk protein concentration. Normalized end-to-end distance (*R^′^_e_*) is defined as the ratio of the distance between the tethered ends (*R_e_*) and the contour length of the DNA (*s*). Bulk protein concentration (*ρ_p_*) is the initial concentration of proteins introduced in the simulation box in units of *µM*. We employ coarse-grained Brownian dynamics simulation to study sequence-dependent protein-DNA condensation. For a detailed discussion of the simulation methods and the different parameters, see the Methods section.

## 3 Results

### 3.1 DNA sequence governs the capillary forces emanating from protein-DNA co-condensates

We seek to uncover the effect of DNA-sequence heterogeneity on the formation of protein-DNA co-condensates and the resultant capillary forces. In particular, we consider two models: homogeneous DNA calling it homo-DNA(Figure. 2A) and heterogeneous DNA calling it hetero-DNA, (Figure. 2B). For a detailed discussion on the choice of parameters, see the methods section, table 1 and 2, and the model section.

First, we keep the normalized end-to-end distance (*R^′^_e_*) fixed and systematically increase the bulk protein concentration (*ρ_p_*). We observe the formation of protein-DNA co-condensate in equilibrium for homogeneous DNA (Figure. 2E). Such co-condensation occurs at a concentration below the saturation concentration for bulk phase separation, which is consistent with previous theoretical [19] and experimental studies [20]. However, it is interesting to note that we observe the formation of a single condensate close to one of the tethered ends, as shown in Figure. 2E (Figure. S1A and S1B). We observed this phenomenon consistently across the various parameters of the model (Figure. S2A). Furthermore, when the protein-protein binding affinity is lower than the protein-DNA binding affinity, elongated condensates emerge (Figure. S2B). As the protein-protein interactions increase, the shape of condensates becomes more spherical. While multiple co-condensates nucleate along the DNA at higher protein concentrations, these condensates fuse and coarsen over time to become a single condensate in equilibrium. The Laplace pressure facilitates this coarsening process, as a single condensate minimizes the surface area compared to multiple smaller condensates in a co-condensate. In contrast, in the case of heterogeneous DNA I, a condensate forms in equilibrium at the high-affinity region of the DNA, as illustrated in Figure. 2F and 2H. While multiple condensates nucleate along the DNA at the beginning of the simulations, the system eventually converges to one condensate, as depicted in the kymograph in Figure. 2J (Figure. S3A and S3B).

**Figure 2:**
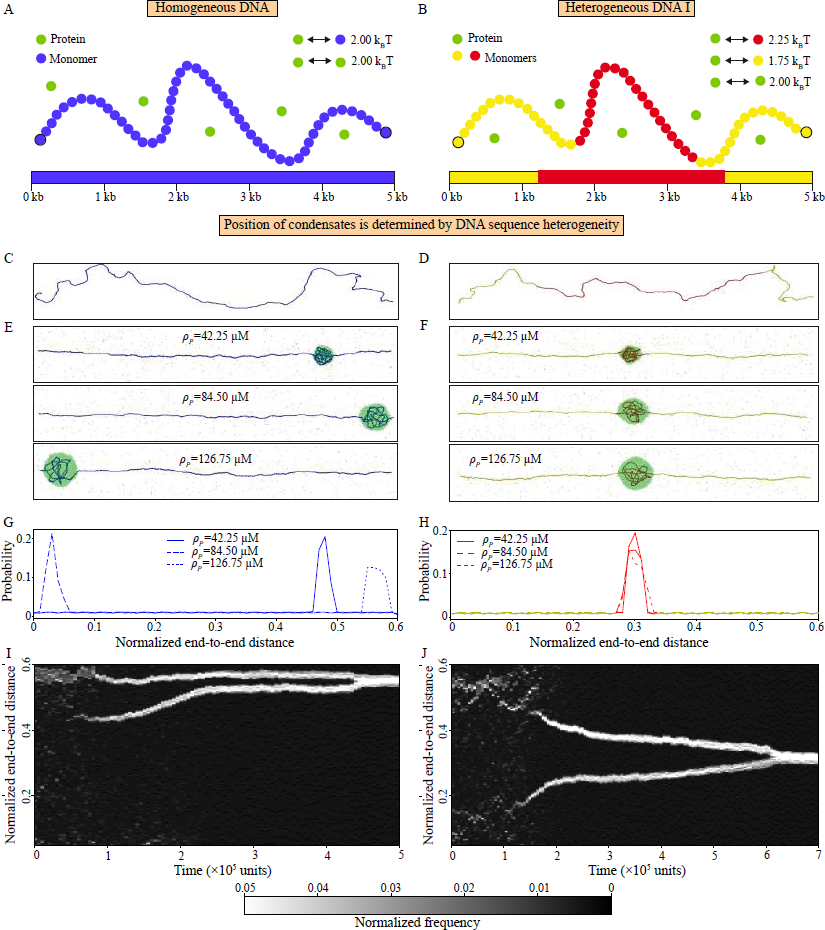
DNA sequence heterogeneity determines position of the condensates. **A** Schematic represents a homogeneous DNA; each monomer (blue) binds to proteins (green) with identical binding affinity (2 *k_B_T*). **B** A heterogeneous DNA model (heterogeneous DNA I) is shown; the model captures the effect of DNA sequence heterogeneity by introducing two types of monomers; monomers (red) at the centre bind proteins with an affinity of 2.25 *k_B_T*, whereas monomers (yellow) at the periphery have an interaction strength of 1.75 *k_B_T*. Proteins bind to each other with an interaction strength of 2.00 *k_B_T*. For parameter details, see table 1 and 2. **C**, **D** Snapshots of DNA configuration for homogeneous and heterogeneous DNA I, in absence of protein, are shown. **E** and **F**, Snapshots of protein-DNA co-condensates in equilibrium for the two models are shown. To generate the plots, we fixed the normalized end-to-end distance (*R^′^_e_* = 0.6) and varied *ρ_p_*. **G**, **H** Probability for each monomer to be inside the condensate (bar width= 5 *σ*) has been shown as a function of position along the DNA for homogeneous (blue) and heterogeneous DNA respectively. **H** The lines shown in yellow and red reflect the DNA with different affinities. **I** and **J**, Representative kymographs for position of condensates along the contour(bar width = 2 *σ*) as a function of time for homogeneous DNA (*R^′^_e_* =0.6, *ρ_p_*= 109.9 *µM*) and heterogeneous DNA I (*R^′^_e_* =0.6, *ρ_p_*= 93.01 *µM*) are shown.

Next, we investigate the effect of protein-DNA condensation on capillary forces as a function of bulk protein concentration (*ρ_p_*) and normalized end-to-end distance of DNA(*R^′^_e_*). For homogeneous DNA, we observe a marginal increase in capillary forces on bare DNA as we increase *ρ_p_* and *_Re_′* (Figure. 3A and 3B). In order to validate that the forces on bare DNA indeed arise from the interface, we calculate total potential energy(see Methods) for each monomer along the contour for homogeneous DNA at *R^′^_e_* = 0.2 and *ρ_p_* = 84.50 *µM* (Figure. S1D). The plot conforms our initial intuition; the potential energy changes sharply for the monomers present at the interface. For heterogeneous DNA I, the capillary force exhibits a non-linear response with respect to *R^′^_e_*. Initially, an increase in the *R^′^_e_* leads to a small increase in capillary force, followed by a sharp jump at *R^′^_e_* = 0.5 (see Figure. 3A). On the other hand, as we increase *ρ_p_*, an opposite trend is observed (Figure. 3B). Capillary force remains constant initially as we increase *ρ_p_*, followed by a gradual decrease after *ρ_p_* = 59.15*µM*. The behavior of capillary forces for heterogeneous DNA I is dictated by the affinity of the interfacial DNA. When *R^′^_e_* exceeds a certain threshold (here 0.5), the higher affinity DNA (shown in red) gets exposed at the interface of the condensate (between compacted DNA and bare DNA), resulting in increased forces exerted on bare DNA (Figure. S3D). In contrast, with increase in *ρ_p_*, condensate volume increases linearly (Figure. S3C), which leads the condensate interface to be exposed to low-affinity (1.75*k_B_T*) DNA (shown in yellow). As a result, there is a decrease in capillary forces exerted on the bare DNA (Figure. 3B and S3E). Overall, our results demonstrate that the protein binding affinity of the interfacial DNA dictates the capillary forces. In the ensuing sections we probe how different kinds of DNA heterogeneity affect capillary forces.

**Figure 3:**
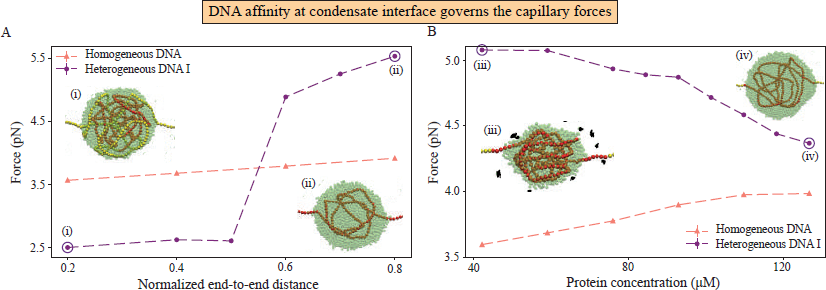
Interfacial DNA affinity governs capillary forces. **A** and **B** Capillary force is plotted as a function of *R^′^_e_* at *ρ_p_*=84.50 *µM* and as a function of *ρ_p_* at *R^′^_e_* =0.6 for homogeneous (salmon) and heterogeneous DNA I (purple) respectively. The snapshots show condensates for heterogeneous DNA I where red represents high affinity monomers while low affinity monomers are shown in yellow. Proteins are shown in green. Snapshots (left to right) show condensates for the following parameter sets: (i) *R^′^_e_* = 0.2 and *ρ_p_*= 84.50 *µM*, (ii) *R^′^_e_* = 0.8 and *ρ_p_*= 84.50 *µM*, (iii) *R^′^_e_* = 0.6 and *ρ_p_*= 42.25 *µM* and (iii) *R^′^_e_* = 0.6 and *ρ_p_*= 126.75 *µM*.

### 3.2 DNA sequence heterogeneity leads to formation of pearl-necklace-like structures

To further explore the effect of DNA heterogeneity on protein-DNA co-condensation, we consider Heterogeneous DNA II, as shown in Figure. 4A (see Model section for more details). Heterogeneous DNA II allows us to explore the impact of a more dispersed heterogeneous DNA sequence on protein-DNA condensation.

Akin to the observations with homogeneous DNA and heterogeneous DNA I (Figure. 2E-2J), with heterogeneous DNA-II, we observe the formation of multiple condensates in the beginning of the simulations. The effect was pronounced at higher protein concentrations(*ρ_p_*) and lower normalized end-to-end distances(*R^′^_e_*). Interestingly, unlike previous DNA models, we find that multiple condensates of different sizes can coexist along the DNA at equilibrium, giving rise to structures reminiscent of pearl necklaces (see Figure. 4B and Figure. S4A). Notably, while a homogeneous DNA leads to a single condensate in equilibrium due to coarsening and fusion, for homogeneous DNA II, the presence of local high affinity regions along the DNA can arrest coarsening. Moreover, the number of condensates varies non-monotonically as a function of *R^′^_e_* andS4C). and *ρ_p_* (see Figure. S4B

In case of bulk phase separation, droplets either fuse or coarsen via diffusion of protein molecules owing to differences in Laplace pressure. Such coarsening leads to the formation of a single condensate in equilibrium. However, in our system, the droplets also interact by applying capillary forces on each other via the DNA, resulting in a tug of war. As we observed in the previous section with the heterogeneous DNA II, such capillary forces depend on the interfacial affinity of the DNA. Evidently, for the multiple condensates to co-exist along the DNA molecule, the system must attain mechanical equilibrium. A systematic examination of droplet volumes and their average interfacial DNA affinity reveals that in scenarios involving condensates of varying sizes, smaller condensates exhibit a greater average interfacial DNA affinity for proteins compared to their larger counterparts (Case 1 and 3 in Figure. 4C). Condensates with similar volumes exhibit similar average interfacial affinity (Case 2 in Figure. 4C). Consequently, as we change *ρ_p_* and *R^′^_e_*, depending on the complex interplay of the position of condensates and their average interfacial affinity, we obtain different number of condensates along the DNA.

Next, we look at the capillary forces as a function of *R^′^_e_* and *ρ_p_*. Capillary forces exhibit a non-monotonic behaviour in both the cases (Figure 4D and 4E) for heterogeneous DNA II. Such non-monotonic behaviour arises due to the block architecture of the DNA based on the average interfacial affinities of DNA (see Method 5.4) for various conditions exhibiting single and multiple condensates at equilibrium. To demonstrate this further, we compute the Pearson correlation coefficient between the capillary forces and the interfacial affinities under various conditions(see Methods 5.5). As shown in Figure 4F, we find a high correlation between them.

The key finding of this section is that DNA sequence heterogeneity can strongly influence the number and position of protein-DNA co-condensates.

**Figure 4:**
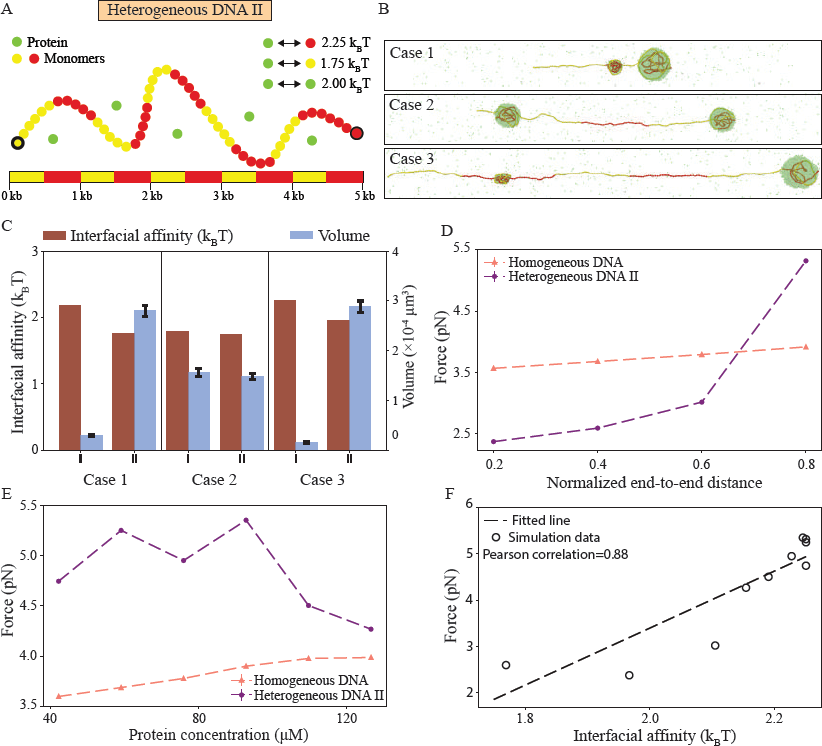
DNA heterogeneity leads to the co-existence of multiple protein-DNA condensates. **A** Schematic represents heterogeneous DNA-II. High affinity monomers (2.25 *k_B_T*, red) are distributed along the contour in form of blocks (length= 50 monomers) interspersed with blocks of low affinity monomers (1.75 *k_B_T*, yellow) of identical length. Proteins (green) interact with each other with interaction strength of 2.00 *k_B_T*. **B**, Snapshots shown for cases with multiple condensates co-existing at equilibrium; the corresponding parameter values are: for Case 1, *R^′^_e_* =0.2 and *ρ_p_*= 84.50 *µM*, for Case 2, *R^′^_e_* =0.4 and *ρ_p_*= 84.50 *µM*, and for Case 3, *R^′^_e_* =0.6 and *ρ_p_*= 84.50 *µM* (top to down). High affinity monomers, low affinity monomers and proteins are shown in red, yellow and green respectively. **C**, Histogram shows average interfacial affinities (maroon) and volumes (blue) for two condensates (I and II) shown in snapshots for all the three cases. **D**,**E** Capillary force is plotted as a function of *R^′^_e_* at *ρ_p_*= 84.50 *µM*, and as a function of *ρ_p_* at *R^′^_e_* =0.6 for heterogeneous DNA II (purple) and homogeneous DNA (salmon) respectively. **F**, Correlation between forces and average interfacial affinities for heterogeneous DNA II has been plotted for all the simulations.

### 3.3 Alterations in the local affinity of interfacial DNA regulate the global capillary forces

To gain a deeper understanding of how DNA sequence impacts forces exerted by protein-DNA cocondensates, next, we study a biologically realistic DNA sequence, the Partial *λ*-DNA (Figure. 5A). For a detailed discussion of this model, see the model section (Table. 3).

Partial *λ*-DNA can lead to the co-existence of multiple protein-DNA co-condensates, similar to the Heterogeneous DNA II model. We find that the number of condensates exhibits a nonmonotonic behavior with respect to *ρ_p_*, and *R^′^_e_*. For simulations in which we observed multiple condensates(Figure. 5B and S5A), we compute the volumes and average interfacial affinities of these condensates (Figure. 5D). In both the instances involving multiple condensates at equilibrium, smaller condensates form interfaces with higher average interfacial affinities when compared to larger condensates. These results are consistent with findings from heterogeneous DNA II model. Notably, findings from both the heterogeneous model II and the partial *λ*-DNA model suggest that to sustain multiple condensates simultaneously along the DNA in equilibrium, smaller condensates must contain interfacial DNA with a higher average affinity.

**Figure 5:**
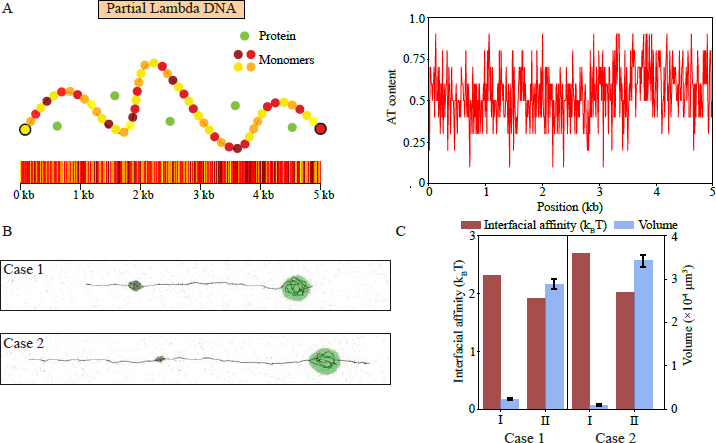
Sequence heterogeneity in partial *λ* DNA engenders multiple condensates at equilibrium. **A**(left), Cartoon representation of heterogeneous DNA III as a last 5 kb of *λ* phage DNA: Eleven kinds of monomers (yellow to maroon) are introduced along the contour based on AT content of 10 bp on partial *λ* DNA sequence. The monomer-protein binding affinities vary in range of 0 to 4 *k_B_T* depending on the AT content of corresponding 10 bp motif that represent the monomer (see the model section and SI). Proteins (green) bind to each other with affinity of 2 *k_B_T*. **A**(right) AT-content of each 10 bp on partial *λ* DNA is shown. **B** Snapshots for the two condensates for the two cases are shown; case 1, *R^′^_e_* =0.4 and *ρ_p_*=84.50 *µM*, and case 2, *R^′^_e_* =0.6 and *ρ_p_*=92.95 *µM* (top to down). **C**, Histogram shows average interfacial affinity (maroon) and volume (blue) for two condensates (I and II) for two cases shown in the snapshots **B**.

The capillary force exhibits a non-monotonic behaviour as a function of *ρ_p_* and *R^′^_e_* (see Figure. 6A and 6B). To probe the non-monotonic behaviour of capillary force, we examine cases associated with the lowest (*ρ_p_*= 59.15 *µM*) and highest (*ρ_p_*= 76.05 *µM*) protein concentrations, as depicted in Figure. 6A. To this end, we analyse the monomer affinities and the probabilities of monomers being at the interface for both the cases(Figure. 6C and 6D). For a change in *ρ_p_* from 59.15*µM* to 76.05*µM*, we observe a 2pN change in capillary force. Although the DNA segments inside the condensate remains nearly identical (snapshots in Figure. 6C and 6D), the sequences at the interface get altered, which lead to significant difference in the capillary forces. A comparison of the interfaces shows that the inclusion of a high-affinity motif at the interface for *ρ_p_*= 76.05 *µM*, leads to pinning of the condensate along the DNA contour, resulting in a smaller interface width (Figure. 6D (bottom panel)). The role of the average affinity of interfacial DNA indeed shows high correlation of 0.91 with capillary forces for different simulations (Figure. 6E). In other words, small changes in affinity of interfacial DNA leads to a significant rise in global capillary forces.

**Figure 6:**
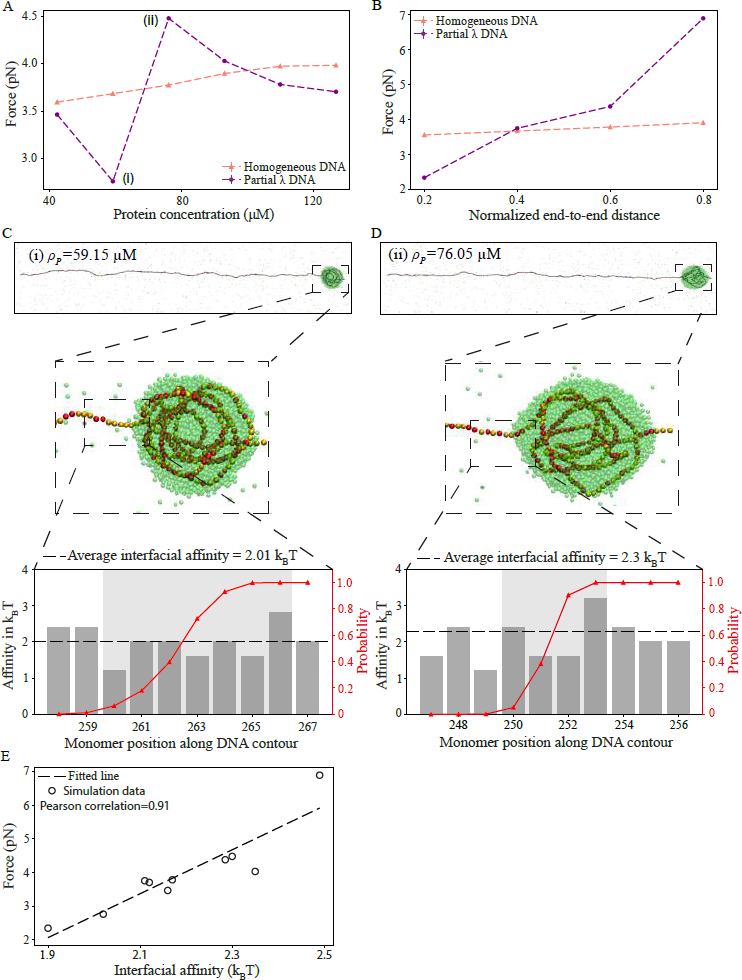
Local interfacial DNA affinity dictates global capillary forces. **A** and **B** represent changes in capillary forces as function of *ρ_p_* and *R^′^_e_* respectively. Non-monotonous behaviour in forces are explained by looking at two conditions (i. *ρ_p_*= 59.15) and (ii. 76.05 *µM*) in **A**. **C**, **D**(top panel), snapshots of condensates are shown for (i) and (ii); the monomers are shown in yellow to red (representing monomer-protein binding affinity in ascending order) while proteins are shown in green. **C**, **D**(bottom panel), bar graphs represent monomer-protein binding affinities; the grey shaded region shows the interface. Probabilities of monomers being inside the condensate are shown in red. **E**, Correlation between capillary forces and average interfacial affinities for all cases with either one or multiple condensates at equilibrium.

### 3.4 HP1 and Sox2 exhibit DNA sequence-dependent co-condensation while Dps shows sequence-independent co-condensation

The results obtained from our simulations can be harnessed to interpret recent experimental results [9,10,20] on protein-DNA co-condensation. . In particular, our results can shed light on the role of DNA sequence on co-condensation. To this end, we analyze the spatial profile and number of co-condensates along the DNA in equilibrium for a set of proteins: nucleoid-associated protein Dps that condenses DNA by bridging its different parts[21], nucleosome binding pioneer transcription factor Sox2[9], and HP1[20] that mediates the formation of heterochromatin.

First, we consider the null homogeneous DNA model. The homogeneous model predicts the formation of a single condensate in equilibrium near one of the tethered ends, as shown in Figure 2. In agreement with this model predictions, a recent study found that Dps forms a single condensate at one of the tethered ends. The concordance between the homogeneous model predictions and experimental data implies that Dps engages in sequence-independent co-condensation with DNA. This finding is consistent with earlier studies that have demonstrated that Dps binds to DNA in sequence-independent manner ([21]. Next, we consider Sox2-DNA co-condensation. In sharp contrast to the predictions of the homogeneous model, the experimental data revealed an opposing trend, with Sox2 forming multiple condensates at various locations along the DNA. These observations falsify the homogeneous model, suggesting that DNA sequence dictates the co-condensation of Sox2. These findings agree with previous studies that have shown DNA sequence-dependent binding of Sox2.

Next, we turn our attention to co-condensation of HP1 with DNA. Hitherto the Binding of HP1 to DNA has been considered as relatively sequence-independent [22]. Based on these previous studies, we hypothesize that the homogeneous DNA model would effectively capture the condensation of HP1 with DNA. Contrary to our expectations, HP1 forms two condensates, with one located near a tethered end and another in the middle of the DNA, thereby falsifying our hypothesis[20]. Evidently, a heterogeneous DNA model with sequence heterogeneity can explain the coexistence of multiple condensates at different locations along the DNA. Our analysis suggests that HP1 exhibits differential binding affinity to DNA.

**Figure 7:**
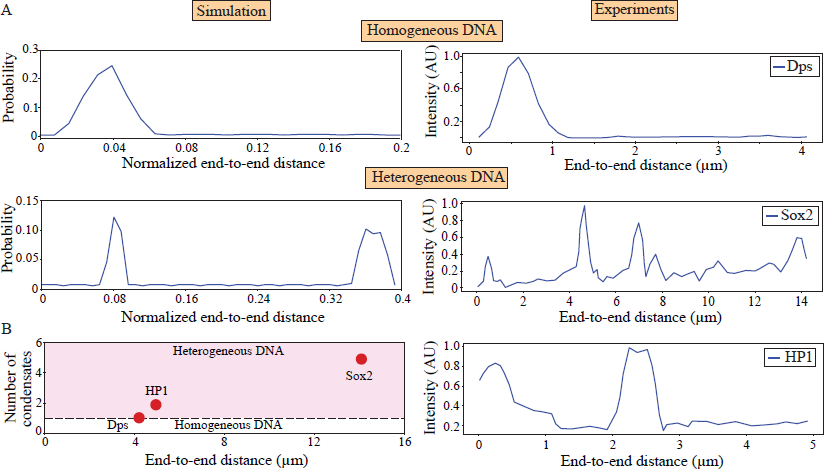
A comparison between simulation and experimental results. **A** Plots on left show Probabilities of monomers being inside the condensate for homogeneous and heterogeneous DNA III respectively(top to bottom). In comparison, on the right, intensity distribution of protein along the DNA has been plotted for Dps (taken from Figure. 4 of [21]), Sox2 (taken from Figure. 1b of [9]) and HP1 (taken from Figure. 4B of [20]) respectively(top to bottom). We extract the data using imageJ and digitizeit (see the Methods section). **B** Scatter plot shows number of condensates(red) as function of end-to-end distance for Dps, Sox2 and HP1, the straight line depicting condensate number of unity reflects sequence-independent co-condensation, whereas region shaded in pink shows sequence-dependent co-condensation of protein and DNA.

## 4 Discussion

Recent evidence suggests that phase separation of chromatin and DNA might play a crucial role in organizing the eukaryotic genome[6,23–29]. However, probing the physical principles of such phase-separated bodies *in vivo* has proved to be challenging. Alternatively, *in vitro* biochemical studies with purified transcription factors (TF) and DNA have shed light on protein-mediated condensation of DNA[8–10]. A series of such *in vitro* studies have shown that protein-protein and protein-DNA interactions lead to condensation of DNA[21,30,31]. However, the role of DNA sequence has thus far remained unclear. Our simulations in this paper provide a framework to decipher the role of DNA sequence on protein-DNA co-condensation.

We find that for a homogeneous DNA, even if multiple condensates form initially, they tend to coarsen over time, eventually resulting in a single condensate—a phenomenon akin to Ostwald ripening. In contrast, multiple co-condensates can co-exist along a heterogeneous DNA, exhibiting pearl-necklace-like structures. For bulk phase separation, droplets interact via diffusion of protein molecules. However, in our model, the droplets also interact by applying capillary forces on each other via the DNA, resulting in a ’tug of war’ effect. The average number and position of various condensates in equilibrium are dictated by a complex interplay of these effects. While recent theoretical studies have explored protein-mediated condensation of DNA, these models primarily focused on the condensation of free DNA polymers[19]. Our simulation results indicate that tethering DNA at both ends and incorporating DNA sequence heterogeneity engender novel physical insights crucial for interpreting existing and future data obtained from *in vitro* studies.

Upon revisiting protein-DNA co-condensation data from previous experimental studies[9,10,20, 21], we find that bacterial nucleoid-associated protein Dps compacts DNA in a sequence-independent manner[21]. In contrast, pioneer factor Sox2 and heterochromatin protein HP1 exhibit sequence-dependent binding to DNA[9,20]. Our findings for Dps are consistent with previous experimental studies. Similarly, Sox2 is known to exhibit high sequence specificity; sox2 recognizes 7-base pair (bp) of (CTTTGTT) DNA sequences. Therefore, our discovery of the co-condensation of Sox2 being influenced by DNA sequence is not surprising. Remarkably, our re-analysis of a recent paper on HP1-mediated DNA condensation revealed the sequence-dependence of HP1 binding[20]. In particular, the existence of multiple condensates, implies that DNA sequence plays a significant role in governing HP1 condensation with DNA. This observation challenges the prevailing understanding of HP1 binding to DNA, which is generally considered to be relatively sequence-independent. Moreover, in the original paper, the authors proposed that the appearance of compacted DNA structures in the center of the DNA molecule is solely driven by large conformational fluctuations occurring in the middle of the molecule[20]. On the contrary, our simulation results for the homopolymer indicate that only a single condensate would form that tends to be located close to one of the two tethered ends. Undoubtedly, additional experiments are needed to test the sequence dependence of HP1 binding to DNA and its role in the condensation of DNA.

Biomolecular condensates in cells form via phase separation[32–36]. Often the formation of these condensates involves various surfaces inside cells such as DNA, microtubule, cytoskeleton, membranes etc[37–39]. The interfaces associated with these structures give rise to interfacial capillary forces. Capillary forces and their physiological implications remain relatively understudied. Our findings suggest that capillary forces arising from the interfacial DNA heterogeneity in protein-DNA co-condensates can generate forces of up to piconewtons. Interestingly, changes in motif level affinity by even one *k_B_T* due to mutations or methylation patterns can lead to two-fold changes in global forces. These forces have the potential to shape the three-dimensional structure of the genome. Moreover, sequence-dependent co-condensation of DNA and proteins can potentially form spatially heterogeneous chromosomal territories. Future studies can shed light on the feasibility of such a mechanism.

Overall, the protein-mediated condensation of DNA is increasingly recognized as a crucial phenomenon for potentially organizing the genome and responsible for other cellular processes[8,10]. Nevertheless, a comprehensive theoretical understanding remains in its early stages. In this light, our modeling framework offers a means to unravel the physical principles governing the sequence-dependent co-condensation of protein and DNA and interpreting experimental observations.

## Acknowledgements

M.G. gratefully acknowledges funding from DBT/Wellcome India Alliance intermediate fellowship (IA/I/21/2/505928). S.C. is supported by the Ramalingaswami Re-entry Fellowship (BT/HRD/35/02/2006), a re-entry scheme of the Department of Biotechnology, Ministry of Science and Technology, Government of India, and Department of Atomic Energy (DAE), Government of India, via Apex project to The Institute of Mathematical Sciences (IMSc), Chennai.

## 5 Methods

### 5.1 Simulation methodology

To simulate the protein-DNA system, we make use of a velocity-Verlet molecular dynamics (MD) scheme, where the temperature is kept *k_B_T* = 1 using a Langevin thermostat

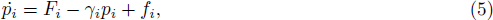

where *F_i_* is force on particle *i* due to interactions with other particles, *f_i_* is the random force generated on particle *i* due to solvent and *γ_i_* is friction coefficient which is kept 0.1 *τ^−^*^1^ in the simulations done. We set integration timestep *δτ* = 0.01 *τ* where *τ* is unit in timescale. The simulations are performed by using the ESPResSo package[40]. Firstly, we generate a linear spring bead model with *N_m_*= 500 in a box of dimensions 20, 20, 600 (*σ* units) and apply force of 0.7 *k_B_T/σ* on first and last monomer in opposite direction to reduce *R^′^_e_*, we simulate the polymer for 1.5 10^6^ *τ* where configurations are stored after every 1000*τ*. Next, we calculate *R^′^_e_* for all the configurations and take the configurations corresponding to *R^′^_e_* values used in simulations. Next, we introduce the respective configurations for different *R^′^_e_* values in a box 80, 80 and 600 (*σ* units) to minimise the polymer interactions with the wall; the ends of the polymer is fixed and persistence length of the polymer is increased from 15 to 50, we simulate the system for 10^6^ timesteps. Next, for equilibration, we reduce persistence length back to 15 and simulate for 3 10^6^ *τ*. For the simulations with proteins, the minimum production time is 3 10^6^*τ* and varies from simulation to simulation. The configurations are stored after every 5000*τ* for analysis, to avoid bias due to temporal correlation.

### 5.2 Model parameters

In this study, the interaction strength for DNA-protein interaction is kept within the range of 0 to 4 *k_B_T* for different DNA models. Indeed, various studies suggest that protein binding affinity to DNA in eukaryotes lies between 0 to 5 *k_B_T* [7]. We set the interaction strength between proteins to be 2 *k_B_T* which is consistent with previous studies [7,41]. We vary the bulk protein concentration (*ρ_p_*) from 42.25 to 126.75 *µM*, which is motivated by an *in vitro* study that studied the co-condensation of HP1 and DNA [20].

### 5.3 Cluster Analysis

In order to identify the condensates in our simulations, we use density based spatial clustering algorithm (DBSCAN)[42] from Scikit-learn. In this algorithm, the variable minimum points is set to 6.0 as the system is 3-dimensional and variable *ε* is calculated from k-distance plot[43]. For *ε* calculation, firstly, distance between particle and its 6th nearest neighbor is calculated for all the particles and arranged in ascending order to plot k-distance plot. Next, to reduce noise in the curve, we fit a third degree polynomial to a window length=199 through Savitzky-Golay filter[44]. Finally, we calculate the point of maximum curvature on the curve and assign the corresponding distance as *ε* in DBSCAN[45]. These calculations were done for 500 configurations spaced over 5000 *τ* from each other.

### 5.4 Force calculation

First, we simulate the polymer in absence of proteins for various normalized end-to-end distances for 3 10^6^*τ* in absence of proteins. Next, we calculate the bond lengths for all 499 bonds and plot its distributions for first and last bonds, as well as other bonds for various end-to-end distances. To this end, we use 500 configurations spaced 10^3^ *τ* from each other. As expected, the distribution of bond length for the tethered ends are identical to the rest of 497 bonds (Figure. S6A). Next, to calculate force on polymer due to stretching in a constant distance ensemble, we use *F_i_* = *K*(*l_i_ − l*_0_) where *K* is the harmonic bond potential, *l_i_* is the bond length of bond *i* and *l*_0_ is the mean bond length calculated for the polymer with free ends. Force is calculated from all the bonds of the polymer and averaged over for each configuration. As before, this is done for 500 configurations spaced 10^3^ *τ* from each other. We then sample 500 lists from the list of forces using bootstrapping to calculate mean of the means and standard deviation of the means for these lists. To express the force in pN, we multiply 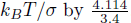. We systematically carry out these calculations for various normalized end-to-end distances and force is plotted as a function of *R^′^_e_* (Figure. S6B). We observe a marginal change in forces up to *R^′^_e_* = 0.8, followed by a sharp rise for *R^′^_e_* = 1 (Figure. S6B), which is consistent with force extension in constant distance ensemble for semi-flexible worm-like chains[46].

For force calculation due to protein-DNA co-condensates, first, the condensates are identified using DBSCAN. Next, we calculate bond length of monomers outside the condensate and calculate force using *F_i_*= *K*(*l_i_ − l*_0_), where *K* is the harmonic bond potential, *l_i_* is the bond length of bond *i*, and *l*_0_ is the mean bond length calculated for the polymer for given *R^′^_e_* mentioned previously. Form each configuration, mean force is calculated and the same analysis is carried out for the last 500 configurations spaced 5 10^3^ *τ* from each other for each case. Procedure of bootstrapping is done (mentioned above) and mean and standard deviation for each case are calculated.

### 5.5 Interfacial affinity

For each configuration, monomer and proteins inside the condensate are detected by DBSCAN. Next, we score 0 or 1 for a monomer based on its absence or presence in the condensate. This is done for all the monomers according to their position along the DNA contour to create an array for each configuration. Next, we average over the arrays generated from the last 500 configurations of the simulations for each monomer to get a probability array. We consider the monomers to be part of interface if the associated probabilities are arranged in increasing (left interface) or decreasing order (right interface) and lie between 0.1 to 1, including the last (left interface) or first (right interface) monomer with probability 1. The reasoning behind this condition is to get all the monomers which form interface in any of the 500 configurations. Next, we calculate the weighted average interfacial affinity *I_a_*, given by

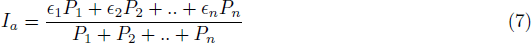

where *ɛ_i_* is the interaction strength between monomer and proteins(*ɛ_MP_*) for monomer *i* and *P_i_* is the probability of monomer *i* inside the condensate. The interfaces which include the tethered monomers are ignored.

### 5.6 Correlation analysis

We calculate the correlation between force and average interfacial affinities using pearson correlation coefficients for heterogeneous DNA II and Partial *λ* DNA models. Mean of the average interfacial affinities of the condensates is taken for correlation calculation.

### 5.7 Volume calculation

To compute the volume of the condensates, we use the maximum distance between the particles belonging to condensate (as detected from DBSCAN is calculated along all three axes. The radius of condensate for each axis is taken as half of the maximum calculated distance along that axis. Volume is then given by

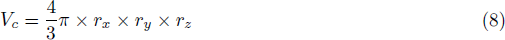

where *V_c_* is the volume of condensate, *r_x_*, *r_y_* and *r_z_* are condensate radii along the three axes. During these calculations, we assign a cutoff of 25 particles in cluster to be considered as a condensate to avoid temporary aggregates in the system. Our results are robust to changes in this cutoff.

### 5.8 Potential energy calculation

We calculate potential energy for each monomer along the contour by calculating their distances with proteins and other monomers. For calculating potential energy contribution from monomer-protein interactions, we use LJ potential employed for monomer-protein interactions in the simulation (see Model details). Similarly, for potential energy contribution from monomer-monomer interactions, we use harmonic and angle potentials as contributions from bonded interactions and LJ potential between monomers as contributions from non-bonded interactions. We sum up all the potential energy contributions for each monomer to get total potential energy. This is done for each monomer over 500 configurations and averaged over for each monomer on the contour(Figure. S1D).

### 5.9 Experimental data analysis

To analyse the experimentally obtained images of protein-DNA co-condensates in various studies, we use imageJ[47]. We extract protein intensity profiles for Sox2 and HP1 by considering the intensity of a line along the DNA. For Sox2, we consider the image in Figure. 1b of [9] and for HP1 we use Figure. 4B of [20]. The peaks in the intensity profiles are counted for estimation of number of condensates. For Dps, we digitize the protein intensity profile from Figure. 4 [21] and plot them as a function of position along the DNA(*µm*)(Figure 7).

## Supplementary Information

### Model details

We use a minimal model to study the protein-DNA co-condensation on end-tethered DNA. We model the DNA as a coarse-grained semiflexible polymer consisting of 500 monomers, and the proteins(*P*) as spherical particles with the same size as the monomers. Each monomer(*M*) in the model maps to 10 base pairs(bps) of DNA, which implies that each protein binds to 10 bps of DNA. Harmonic springs link the adjacent monomers, whereas harmonic angle potential between consecutive bonds models the semi-flexibility of DNA. The bond length potential is given by,

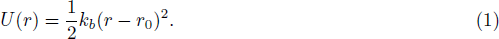

Here, *k_b_* and *r*_0_ are the bond length energy and equilibrium bond length, respectively. We set *k_b_* = 100 *k_B_Tσ^−^*^2^ and *r*_0_ = *σ*. The bond angle potential is given by,

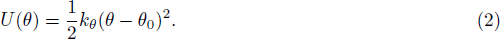

Here, *k_θ_* and *θ*_0_ are the bond angle energy and equilibrium bond angle. We set *k_θ_* = 15 *k_B_T* and *θ*_0_ = 180°. *k_θ_* relates to the persistence length(*l_p_*) of DNA as,

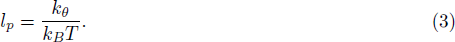

We consider four different DNA sequences in the study. In each of the sequences, DNA is modeled as a self-avoiding polymer with only repulsive interaction between the monomers. To model the repulsion between the monomers, we use Weeks-Chandler-Anderson (WCA) potential[48], which is a purely repulsive form of Lennard-Jones potential. To model the non-bonded interactions among monomer-protein and protein-protein, we use the LJ potential with attractive tail.

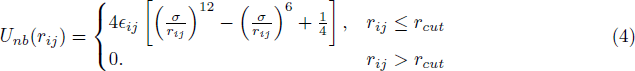

where *r_ij_* is the distance among the particles *i* and *j*.

However, the details of the monomer-protein and protein-protein interactions vary for different sequences. For the homogeneous DNA (Figure 2A) which consists of only one type of monomers (A), we have three interacting pairs, namely, AA, AP and PP. As stated above, we keep purely repulsive interaction between AA pairs to model the volume exclusion of monomers. To begin with, we keep the strength of monomer-protein attraction (*ɛ_AP_*) and as well as protein-protein attraction (*ɛ_P P_*) equals to 2 *k_B_T*. Table1summarizes the non-bonded interaction parameters for homogeneous DNA-protein co-condensate system.

For the heterogeneous DNA I (Figure 2B) and heterogeneous DNA II (Figure 3A), we consider the DNA to be made up of two types of monomers, A and B. A monomers attract the proteins P with 1.75 *k_B_T* strength, whereas B monomers attract the proteins P with 2.25 *k_B_T* strength. In both the sequences, we keep equal number (250 each) of A and B monomers, such that mean DNA-protein affinity is 2 *k_B_T*, which is same as that for the homogeneous DNA. However, the two heterogeneous DNA models differ in terms of the sequence heterogeneity. Heterogeneous DNA I (Figure 2B) has a -AAAAA-BBBBB-AAAAA-like architecture with the central 250 B monomers having stronger affinity for the proteins P. The central B region is franked by two stretches of A monomers each of length 125, having weaker affinity for the proteins P. Heterogeneous DNA II (Figure. 3A) has a block copolymer(-AABBAAABBAABB-) like architecture where 50 monomers of A are followed by 50 monomers of B. The interaction parameters for heterogeneous model I and II are summarized in Table 2.

**Table 1:** Non-bonded interaction parameters for all the particle pairs in the homogeneous DNA-protein cocondensate system: A represent the monomers, and P represents the proteins.

| $ij$ | $\epsilon_{ij} (k_B T)$ | $r_{cut} (\sigma)$ |
| --- | --- | --- |
| AA | 1.00 | $2^{\frac{1}{6}}$ |
| AP | 2.00 | 2.5 |
| PP | 2.00 | 2.5 |

**Table 2:** Non-bonded interaction parameters for all particle pairs. A, and B are monomers and P represents the proteins. Monomer A has weaker affinity for the proteins (1.75 *k_B_T*) and monomer B has stronger affinity for the proteins (2.25 *k_B_T*).

| $ij$ | $\epsilon_{ij} (k_B T)$ | $r_{cut} (\sigma)$ |
| --- | --- | --- |
| AA | 1.0 | $2^{\frac{1}{6}}$ |
| AB | 1.0 | $2^{\frac{1}{6}}$ |
| BB | 1.0 | $2^{\frac{1}{6}}$ |
| AP | 1.75 | 2.5 |
| BP | 2.25 | 2.5 |
| PP | 2.0 | 2.5 |

Our fourth DNA sequence is the last 5 Kbp stretch of *λ*-phage DNA (partial *λ* DNA) sequence. We assume that the protein P binds to the AT rich region of the sequence. Since one monomer in our model corresponds to 10 bps, each monomer may have different affinity for the protein P, based on the AT content of underlying 10 bp sequence. If the all the 10 bps are either Adenine or Thyamine nucleotide, then it has 100% AT content, whereas if none of the sequences is A or T, then it has 0% AT content. So, the AT content of a 10 bps sequence can only have 11 possible values. For this purpose, we consider our system to be made up of 11 types of monomers (A, B, C, D, E, F, G, H, I, J, K), each corresponding to a different AT content of the 10 bps sequence. Using this formalism, we find the AT content of the partial *λ* DNA as a function of the sequence length (Figure. 5B). Next, we assign different monomer-protein affinities to these 11 monomers on a scale from 0 to 4 *k_B_T*. We summarize the monomer-protein interaction parameters in Table (3). We keep *ɛ_P P_* = 2 *k_B_T* as in the previous sequences.

**Table 3:** AT content of 10 bps of eleven kinds of monomers in the partial *λ* DNA model, and their interaction affinities with protein P.

| AT content(%) | monomer (M) | $\epsilon_{MP} (k_B T)$ |
| --- | --- | --- |
| 0 | A | 0.0 |
| 10 | B | 0.4 |
| 20 | C | 0.8 |
| 30 | D | 1.2 |
| 40 | E | 1.6 |
| 50 | F | 2.0 |
| 60 | G | 2.4 |
| 70 | H | 2.8 |
| 80 | I | 3.2 |
| 90 | J | 3.6 |
| 100 | K | 4.0 |

### 5.10 Supplementary figures

**Figure S1:**
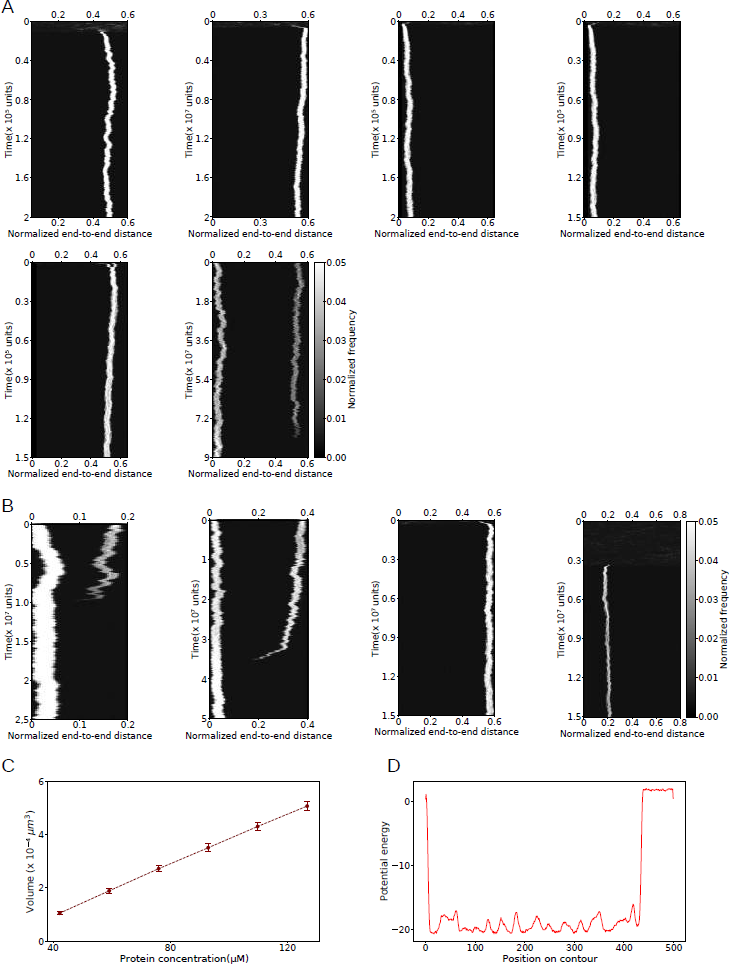
A homogeneous DNA co-condense with proteins to form a single condensate close to one of the tethered ends at equilibrium. A Representative kymographs. DNA density distribution along the contour (bar width = 2 *σ*) as a function of time for homogeneous DNA at different *ρ_p_*= 42.25, 59.15, 76.05, 92.95, 109.85 and 126.75 (in *µM*) (left to right) respectively for constant *R^′^_e_* = 0.6. **B** Representative kymographs for homogeneous DNA at different *R^′^_e_* = 0.2, 0.4, 0.6 and 0.8 (left to right) respectively for *ρ_p_*= 84.50 *µM*. **C** Volume of the condensate is plotted as a function of *ρ_p_*. **D** Total potential energy is shown for each monomer along the contour for homogeneous DNA at *R^′^_e_* =0.2 and *ρ_p_*= 84.50 *µM* (see Methods for potential energy calculations).

**Figure S2:**
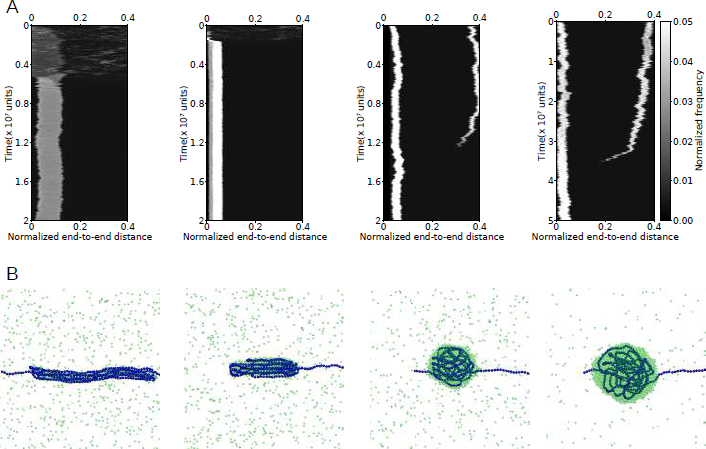
Co-condensation with homogeneous DNA leads to single condensate at various protein-protein interaction strengths. A, Representative kymographs for position of condensates along the contour(bar width = 2 *σ*) as a function of time for homogeneous DNA (*R^′^_e_* =0.5, *ρ_p_*= 85.50 *µM*) at *ɛ_PP_* ; 1, 1.5, 1.8 and 2 *k_B_T* (left to right) respectively. B, Snapshots (i), (ii), (iii), and (iv) represent condensates at *ɛ_P P_* = 1, 1.5, 1.8 and 2 *k_B_T* (left to right) respectively. Homogeneous DNA monomers are shown in blue and proteins are shown in green.

**Figure S3:**
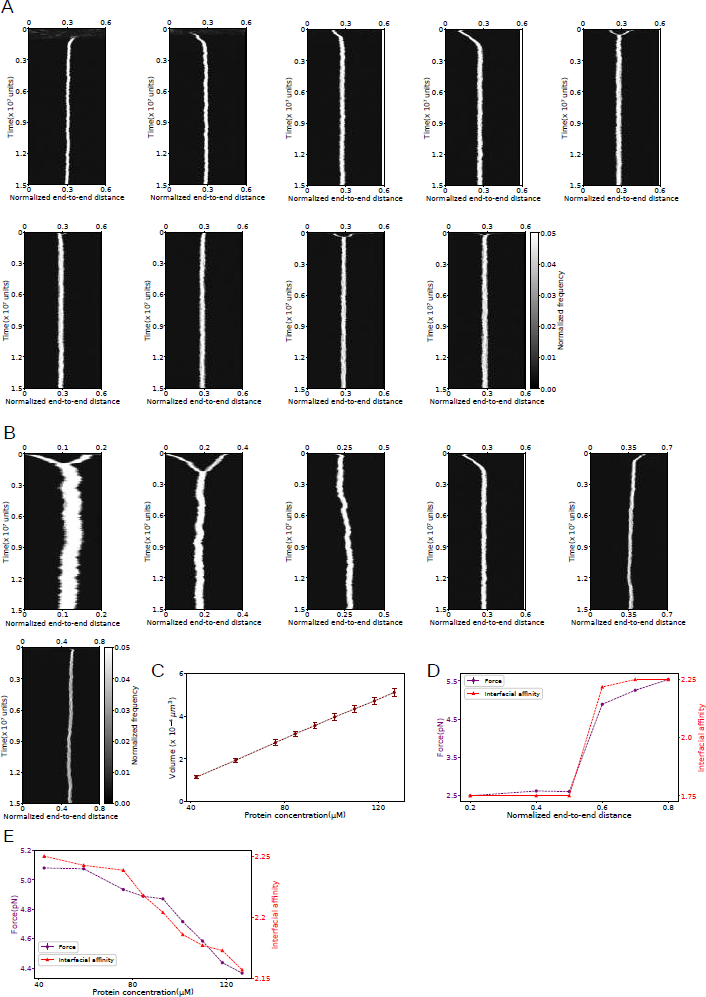
Condensates positions are governed by DNA sequence heterogeneity. A Representative kymographs for DNA density distribution along the contour (bar width = 2 *σ*) as a function of time for heterogeneous DNA I at different *ρ_p_*= 42.25, 59.15, 76.05, 92.95, 101.40, 109.85, 118.30 and 126.75 (in *µM*) (left to right) respectively for *R^′^_e_* = 0.6. **B** Representative kymographs for heterogeneous DNA at different *R^′^_e_* = 0.2, 0.4, 0.5, 0.6, 0.7 and 0.8 respectively for *ρ_p_*= 84.50 *µM*. **C** Volume of the condensate is plotted as a function of *ρ_p_* for heterogeneous DNA I (see Methods for volume calculations). **D** Force (purple) and average interfacial affinity (red) as a function of *R^′^_e_* at *ρ_p_*=84.50 *µM*. **E** Force (purple) and average interfacial affinity (red) is plotted as a function of *ρ_p_* at *R^′^_e_* = 0.6 for heterogeneous DNA I. **F** Total potential energy for each monomer is plotted along the contour for heterogeneous DNA I at *R^′^_e_* = 0.2 and *ρ_p_*= 84.50 *µM*.

**Figure S4:**
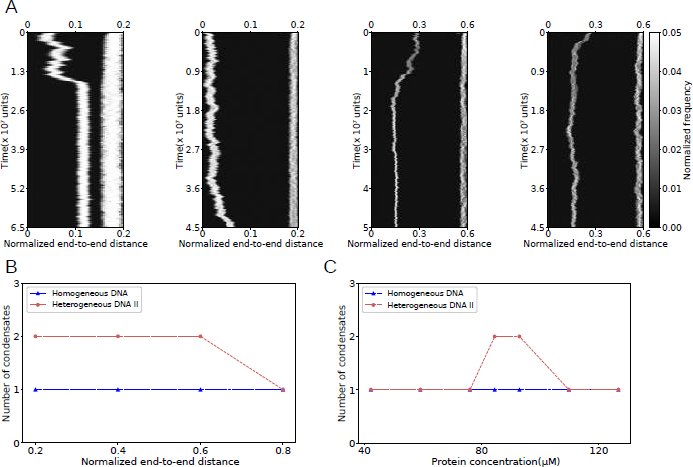
DNA heterogeneity leads to co-existence of multiple co-condensates at equilibrium. A Representative kymographs of DNA density distribution along the contour (bar width = 2 *σ*) as a function of time for heterogeneous DNA II for 4 different cases that lead to multiple condensates at equilibrium; case 1 *R^′^_e_* = 0.2 and *ρ_p_*= 84.50 *µM*, case 2 *R^′^_e_* = 0.4 and *ρ_p_*= 84.50 *µM*, case 3 *R^′^_e_* = 0.6 and *ρ_p_*= 84.50 *µM*, case 4 *R^′^_e_* = 0.6 and *ρ_p_*= 92.95 *µM* (left to right). **B** Number of condensates at equilibrium is plotted as a function of *R^′^_e_* at *ρ_p_*= 84.50 *µM* for homogeneous DNA (shown in blue) and heterogeneous DNA II (shown in maroon). **C** Number of condensates at equilibrium is plotted as a function of *ρ_p_* at *R^′^_e_* heterogeneous DNA II (shown in maroon). = 0.6 for homogeneous DNA (shown in blue) and heterogeneous DNA II (shown in maroon).

**Figure S5:**
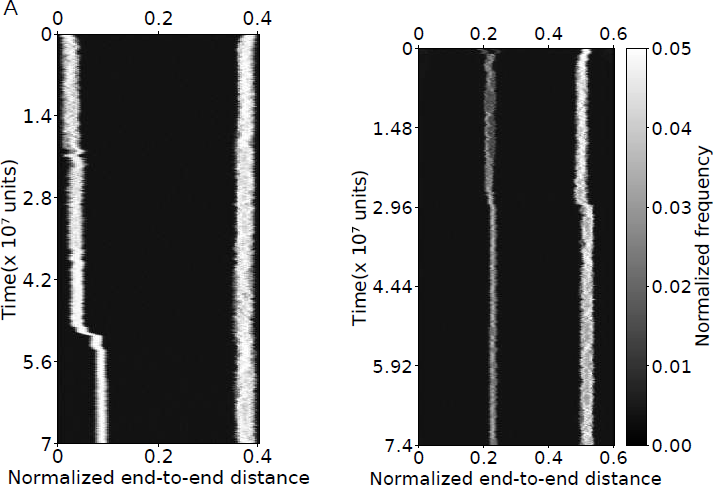
Partial *λ* DNA can sustain multiple condensates. **A** Representative kymographs of DNA density distribution along the contour (bar width = 2 *σ*) as a function of time for heterogeneous DNA III (partial *λ* DNA) for two cases which lead to multiple condensates at equilibrium; case 1 *R^′^_e_* = 0.4 and *ρ_p_*= 84.50 *µM*, case 2 *R^′^_e_* = 0.6 and *ρ_p_*= 92.95 *µM* (left to right).

**Figure S6:**
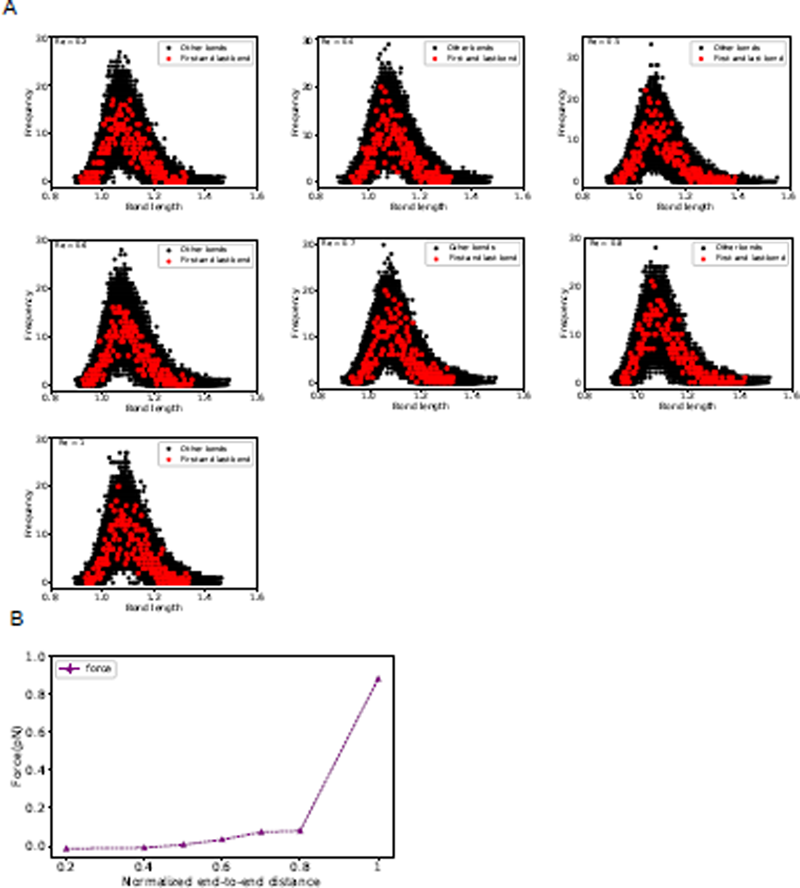
Bond length distribution and force extension curve for the semiflexible polymer. **A** Bond length distributions for first and last bond (shown in red) and other bonds (shown in grey) for the polymer has been plotted for different normalized end-to-end distances *R^′^_e_* = 0.2, 0.4, 0.5, 0.6, 0.7, 0.8 and 1 in absence of proteins. **B** Force calculated from all bonds is plotted as a function of normalized end-to-end distance for the semiflexible polymer (see Methods for force calculations).

## Notes

### Competing Interest Statement

The authors have declared no competing interest.

